# Cortical tracking of prediction error during perception of connected speech

**DOI:** 10.1101/2025.07.18.665498

**Authors:** James M. Webb, Ediz Sohoglu

## Abstract

Prediction facilitates speech comprehension, but how predictions are combined with sensory input during perception remains unclear. Prior studies suggest that prediction error computations — which represent the difference between heard and expected speech sounds — play a central role in speech perception. However, these studies have relied on non-ecological listening conditions, involving isolated words predicted by artificial cues. In two experiments, we presented 61 listeners with semantically coherent sentences that ended in words of varying predictability. We also manipulated the signal quality of the final word by applying time-limited degradation. Using linear encoding models of EEG responses, we found that neural representations of speech features were jointly influenced by top-down predictions and bottom-up signal quality. Specifically, for unpredicted final words, increasing signal quality resulted in enhanced neural tracking of speech acoustic modulations in low frequency EEG (0-2 Hz delta band). In contrast, for strongly predicted final words, greater signal quality led to reduced tracking of speech modulations. Computational simulations revealed that this interaction is consistent with prediction error computations, but not with an alternative signal sharpening model. These findings extend the evidence for prediction error computations to more naturalistic listening situations.

## Introduction

Because of the inherent ambiguity of naturalistic speech, listeners make use of prior knowledge during speech perception. Prior knowledge encompasses not only linguistic knowledge but also situational context and knowledge about the world. Understanding how listeners integrate prior knowledge with sensory input is crucial for understanding the mechanisms underlying speech perception.

It has been suggested that predictions are combined with the bottom-up speech signal in a process of Bayesian inference [1–3]. Under Bayesian inference, speech perception is a succession of inferences that combine prior knowledge with incoming sensory data as the perceptual system attempts to find the most likely interpretation of the speech signal. In doing so, the listener continuously adjusts and refines their predictions given new sensory evidence.

Here we build on previous work seeking to establish how Bayesian inference is implemented neurally. One possibility is that neural representations reflect a weighted average of the sensory input and prior expectations [4]. Under this account, more accurate predictions and higher signal quality should both act to enhance or ‘sharpen’ neural representations, just as they both act to enhance perceptual clarity. Alternative models propose that sensory features are suppressed by predictions, with unexpected features (prediction errors) prioritised for processing and updating predictions [5–8]. In predictive coding models, sharpened signal representations and prediction errors are dually encoded in distinct neural populations [9,10].

A brain measure that has widely been used to investigate predictive processing is the N400 component. It is characterised by a negative deflection occurring 200-600 ms after word onset and is particularly strong when a word is unexpected given the prior context (e.g. sentence or word prime [11,12]. Accordingly, the N400 is reduced in amplitude for predictable words. This observation, and other phenomena characterised by reduced neural responses to predicted stimuli [13–15] is commonly attributed to the minimisation of prediction error. However, reduced activity is equally compatible with sharpened responses because, under this account, neuronal activity representing competing features (i.e. ‘noise’) is suppressed [16–19]. For the same reason, more recent demonstrations of neural responses correlating with speech predictability during naturalistic (e.g. story) listening [20–24] also do not conclusively support prediction error over sharpened signal accounts.

It has been suggested that a more effective way to distinguish between accounts is by using information-based analysis methods [17]. Rather than measuring the overall strength of neural responses irrespective of speech content, information-based methods permit the analysis of responses to specific speech features. The finer-grained resolution of these methods potentially provides the opportunity to better distinguish sharpened signals from prediction errors. Indeed, using computational simulations, Blank and Davis [16] demonstrate that the information content in simulated prediction errors is interactively modulated by prior expectations and signal quality. This is because when predictions are accurate, prediction error representations are more greatly suppressed as signal quality increases. Yet when predictions are uninformative, increasing signal quality leads instead to enhanced prediction errors as there is more information left ‘unexplained’. Importantly, a very different result is observed when simulating sharpened signals. In this case, the information content in simulated patterns is enhanced by prior expectations and signal quality in an additive fashion, in much the same way as perceptual outcomes.

Blank and Davis [16] went on to observe an interactive effect of prior expectation and signal quality on multivoxel patterns in the superior temporal lobe, exactly as observed when simulating prediction errors. Using MEG and linear encoding models, Sohoglu and Davis [25] showed that this interaction is also apparent when examining the extent to which the cortex ‘tracks’ speech acoustic features.

The information-based neural findings above therefore suggest that brain responses encode prediction errors, which imposes constraints on the underlying neural computations. They provide direct support for predictive coding models and argue against models that only propose sharpened signal computations. However, in this previous work, listeners heard single words while prior knowledge was manipulated by presenting matching or nonmatching text before the speech. This is far removed from real-world conditions where listeners need to combine predictions with sensory input continuously (i.e. during perception of connected speech) and often without the use of external cues. As such, these findings are susceptible to the criticism that the prediction effects which occurred were encouraged by the experimental design and may not be representative of naturalistic perception [22,26].

In the two experiments reported here, listeners heard sentences in which strong or weak predictions for the final word could be obtained from the preceding speech context without external cues (see Figure 1A and 1B). To manipulate final word signal quality independently of listeners’ predictions, we developed a novel form of noise-vocoding that allowed us to vary speech signal quality in a time-limited fashion i.e. separately during the context and final word periods.

**Figure 1.**
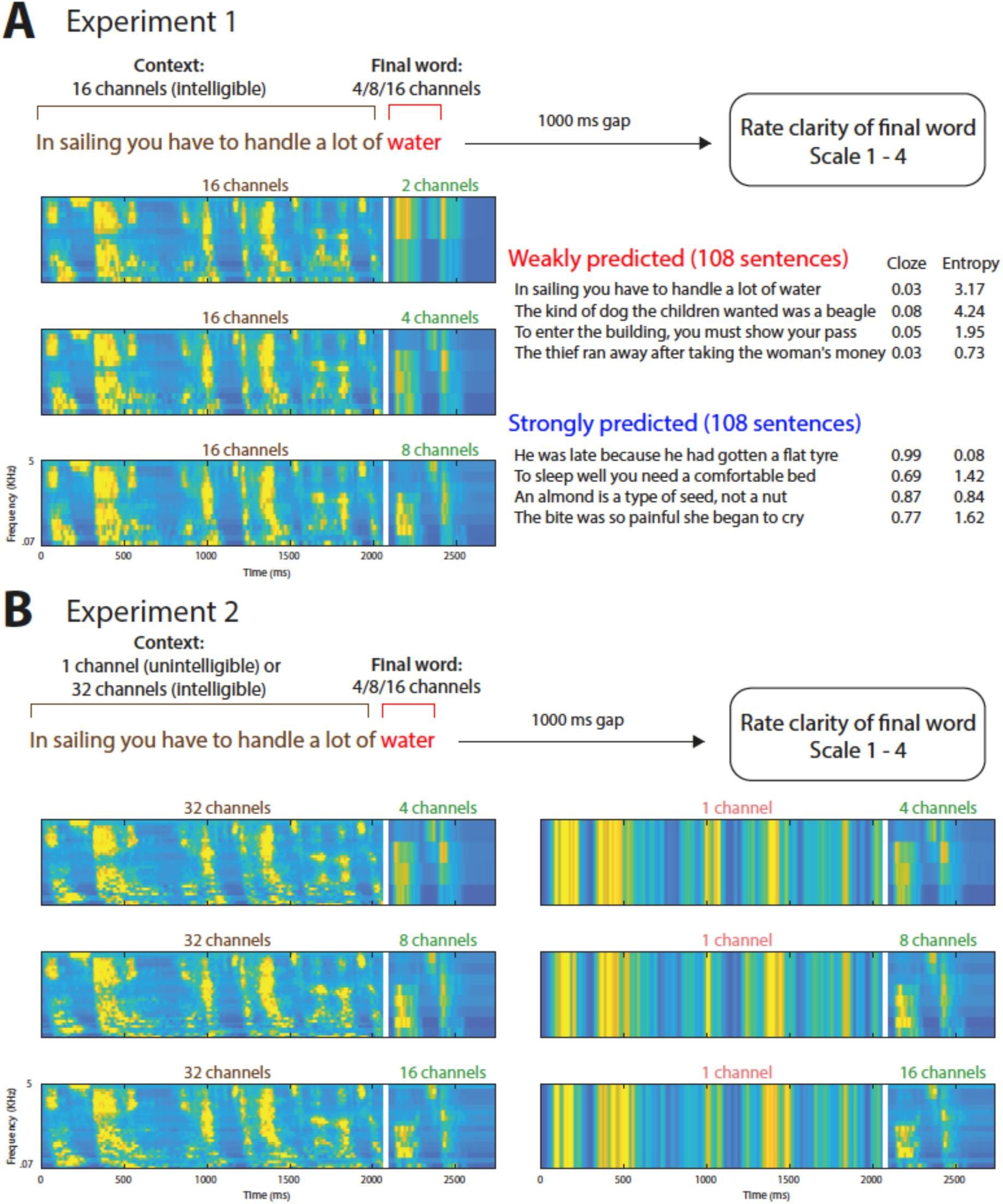
Procedure. **A)** For Experiment 1, on each trial listeners heard a sentence with a weakly or strongly predicted final word. The final word was noise-vocoded using 2, 4, or 8 channels. The preceding sentence context was always highly intelligible (16 channel vocoded). Participants were required to rate the clarity of the final word in each sentence on a scale of 1 – 4. Example sentences are depicted along with final word cloze probabilities and context entropy. Time-frequency plots represent auditory spectrograms [28] for the sentence “In sailing you have to handle a lot of water”. The vertical white line marks the onset of the final word (for visualisation purposes only as no silence was inserted between the context and final word). **B)** For Experiment 2, the final word was noise-vocoded using 4, 8, or 16 channels. The preceding sentence context was noise-vocoded using either 32 channels (highly intelligible) or 1 channel (unintelligible).

To examine how cortical tracking of speech is modulated by signal quality and prior expectation, we regressed acoustic speech features against 64 channel EEG using linear encoding models [27]. In a sharpened signal account, increasing signal quality or prediction strength for the final word should result in improved tracking of the heard word in neural responses (Figure 2C). Whereas in a prediction error account, there should be an interaction effect such that increasing signal quality results in better tracking of the heard word when final word predictions are weak and the opposite when predictions are strong (Figure 2D).

**Figure 2.**
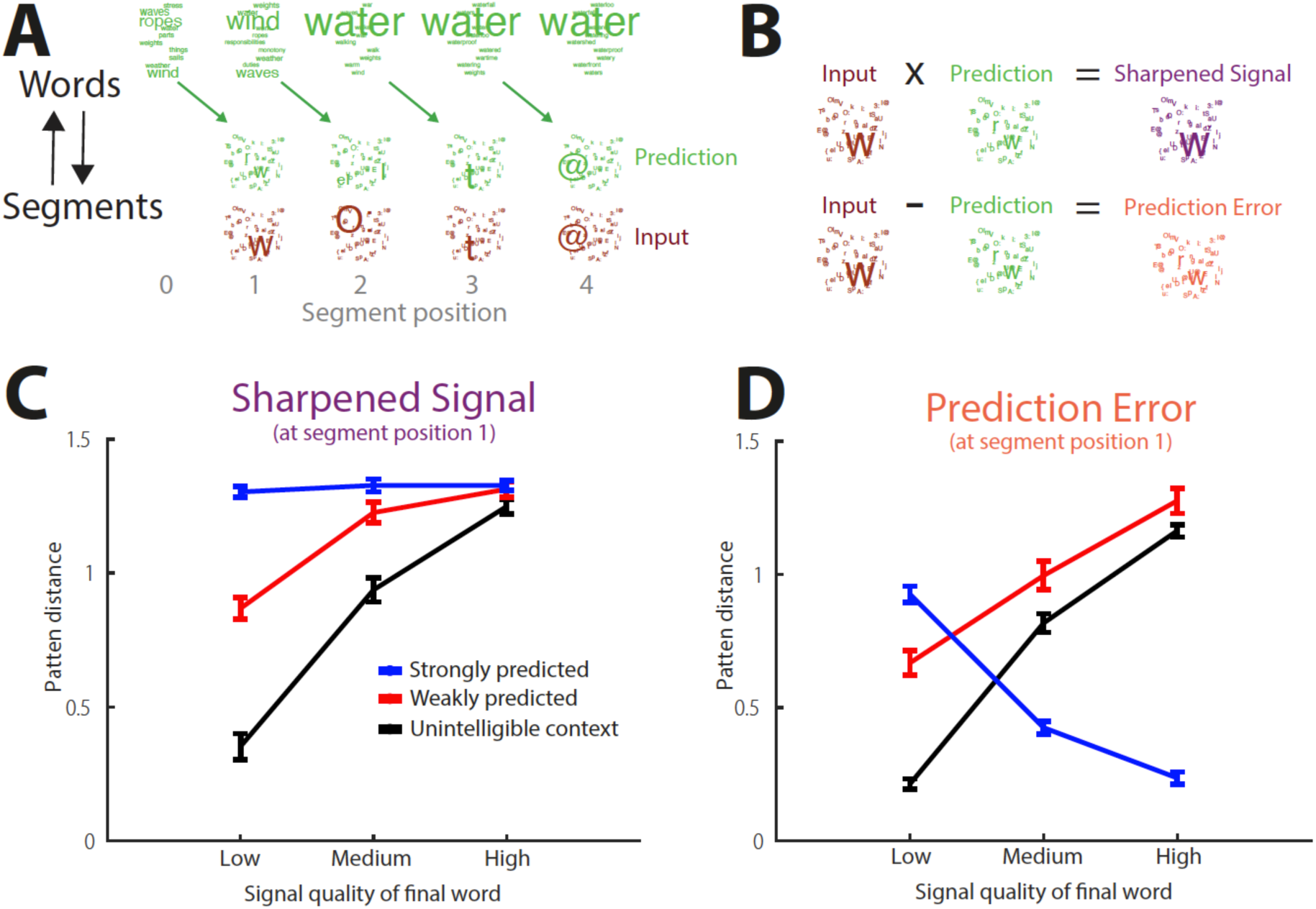
Computational simulations. **A)** Model representations for the weakly predicted final word (/w O: t @/) in the sentence “In sailing you have to handle a lot of water”, simulated in a high signal quality condition. Model representations are visualised using word and segment clouds with larger text indicating higher probabilities and segment labels denoting phonetic transcriptions in SAMPA format. (Top row) Word level representations show posterior probabilities as each segment in the input is heard (only the ten words with the highest probabilities at any one timepoint are shown for illustration purposes). Word posteriors provide predictions for which segment will be heard next (second row, segment predictions shown in green). Segment predictions are then combined with segment input probabilities (‘Input’, third row; shown in brown). **B)** Segment predictions and input representations are combined through sharpened signal (multiplication) or prediction error (subtraction) computations with outcomes depicted as purple and orange segment clouds, respectively (for the onset segment of water). **C)** Mean pattern distances at final word onset (segment position 1) for the sharpened signal simulations. We simulated Experiment 2 which comprised a more complete set of experimental conditions (including the unintelligible context condition). Error bars represent the standard deviation. **D)** Same as panel C but for the absolute prediction error.

## Results

### Overview of experiments

In two experiments, listeners heard sentences in which strong or weak predictions for the final word could be obtained from the preceding sentence context. To manipulate final word signal quality independently of listeners’ predictions, we used noise-vocoding to vary spectral detail separately during the context and final word periods (see Figure 1A and 1B for a depiction of these experimental manipulations; see Figure 3 for a depiction of stimulus properties).

**Figure 3.**
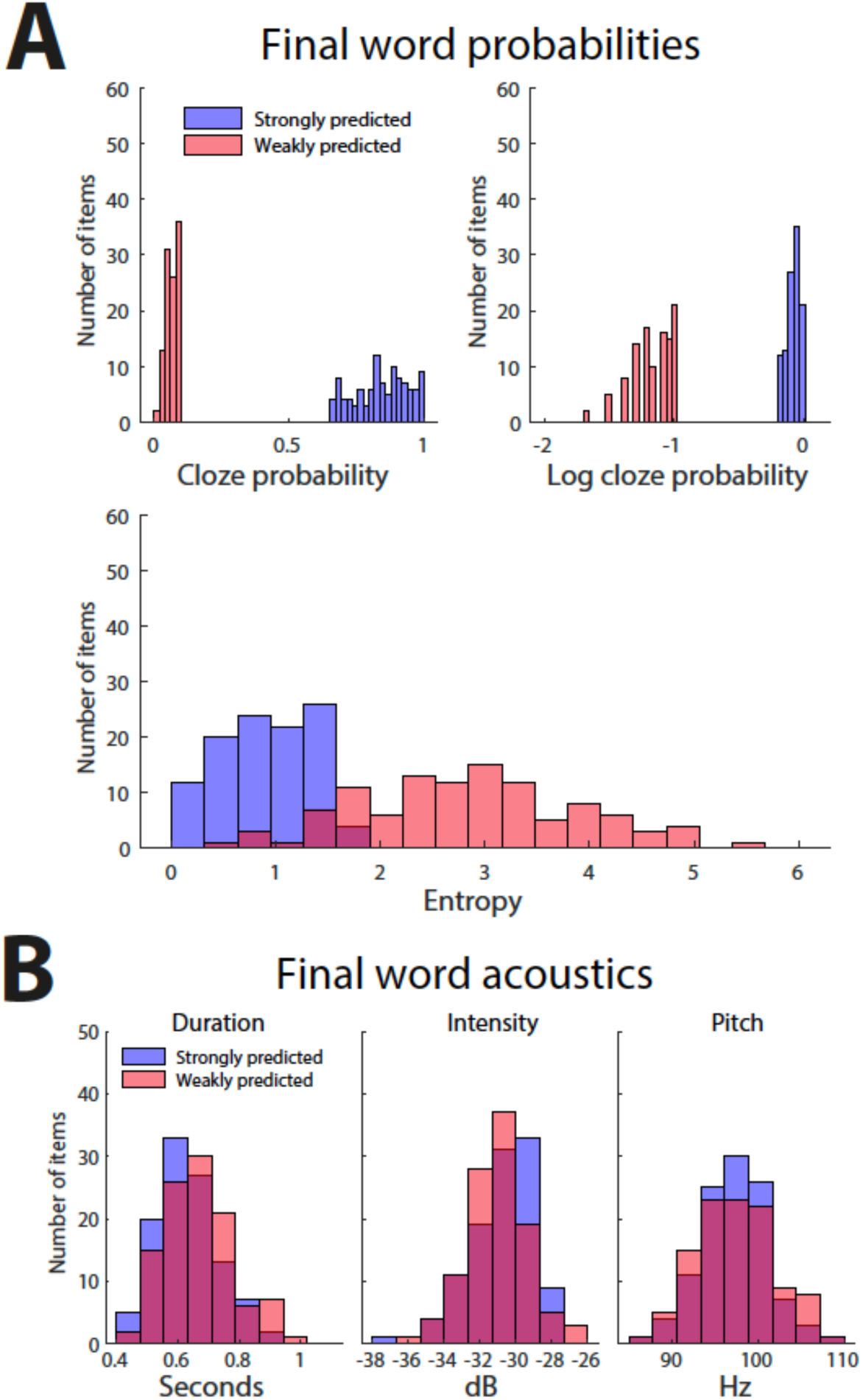
Stimulus properties. **A)** Strongly predicted final words had higher cloze probabilities than weakly predicted final words (upper graphs). In the strongly predicted condition, sentence contexts were also more constraining on average (i.e. had lower entropy) than in the weakly predicted condition. **B)** There were no reliable differences in final word duration, intensity, or pitch.

In Experiment 1 (Figure 1A), the sentence context portion was always highly intelligible (16 channel vocoded) but the final word varied between 2, 4 and 8 channels. In Experiment 2 (Figure 1B), we presented sentences in which the final word was noise-vocoded with 4, 8 or 16 spectral channels, enabling a more extreme manipulation of signal quality. In Experiment 2, we also presented the context portion of the sentences with either 32 spectral channels (highly intelligible) or 1 channel (unintelligible). The latter condition was presented so that we could observe neural responses when listeners are unable to make predictions for the final word.

### Computational simulations: At word onset

To confirm that our experimental paradigm can distinguish sharpened signals from prediction errors, we performed simulations of sharpened signal and prediction error computations. Both sharpened signal and prediction error computations are derived from posterior word probabilities, as estimated using Bayes theorem (see Methods). We applied this model to the stimuli of Experiment 2 where signal quality and prediction strength varied more extremely than in Experiment 1.

Example model outputs are shown in Figure 2A and 2B for the final word in the weakly predicted sentence “In sailing you have to handle a lot of **water**”. To summarise the results of these simulations, we quantified how strongly speech content was represented in simulated sharpened signals (Figure 2C) and prediction errors (Figure 2D). We achieved this by computing the dissimilarity between model representations (Euclidean pattern distances) for all pairs of items within each condition. We focus on the model’s responses at final word onset (segment position 1) as this is the earliest position at which predictions can be combined with the speech input. In a subsequent section, we will present the model’s responses at later segment positions.

As shown in Figure 2C and 2D, whereas final word prediction strength and signal quality both act to enhance sharpened signals, these manipulations have opposite effects on prediction errors. The opposing effects on prediction errors take the form of a crossover interaction. In particular, increasing signal quality results in increased prediction error when the final word is unpredicted (unintelligible context and weakly predicted conditions). The opposite effect – reduced prediction error with increasing signal quality – occurs when the final word is strongly predicted. These simulations therefore confirm that our experimental manipulations should produce different outcomes if neural responses reflect sharpened signals versus prediction errors.

### Experiment 1

#### Behaviour

In Experiment 1, the sentence context portion was always highly intelligible (16 channel vocoded) but the final word varied between 2, 4 and 8 channels (see Figure 1A). As shown in Figure 4A, final word clarity ratings were enhanced not only by final word signal quality (F(1.32, 38.37) = 752.42, p < 0.001) but also by final word prediction strength (F(1, 29) = 83.42, p < 0.001). There was also an interaction between final word prediction strength and signal quality (F(2, 58) = 9.68, p < 0.001). Visual inspection of the data in Figure 4A suggests that this interaction reflects an increased influence of prediction strength in the 2 channel and 4 channel conditions, as compared with the 8 channel condition. These results confirm that both signal quality and prediction strength enhance speech clarity.

**Figure 4.**
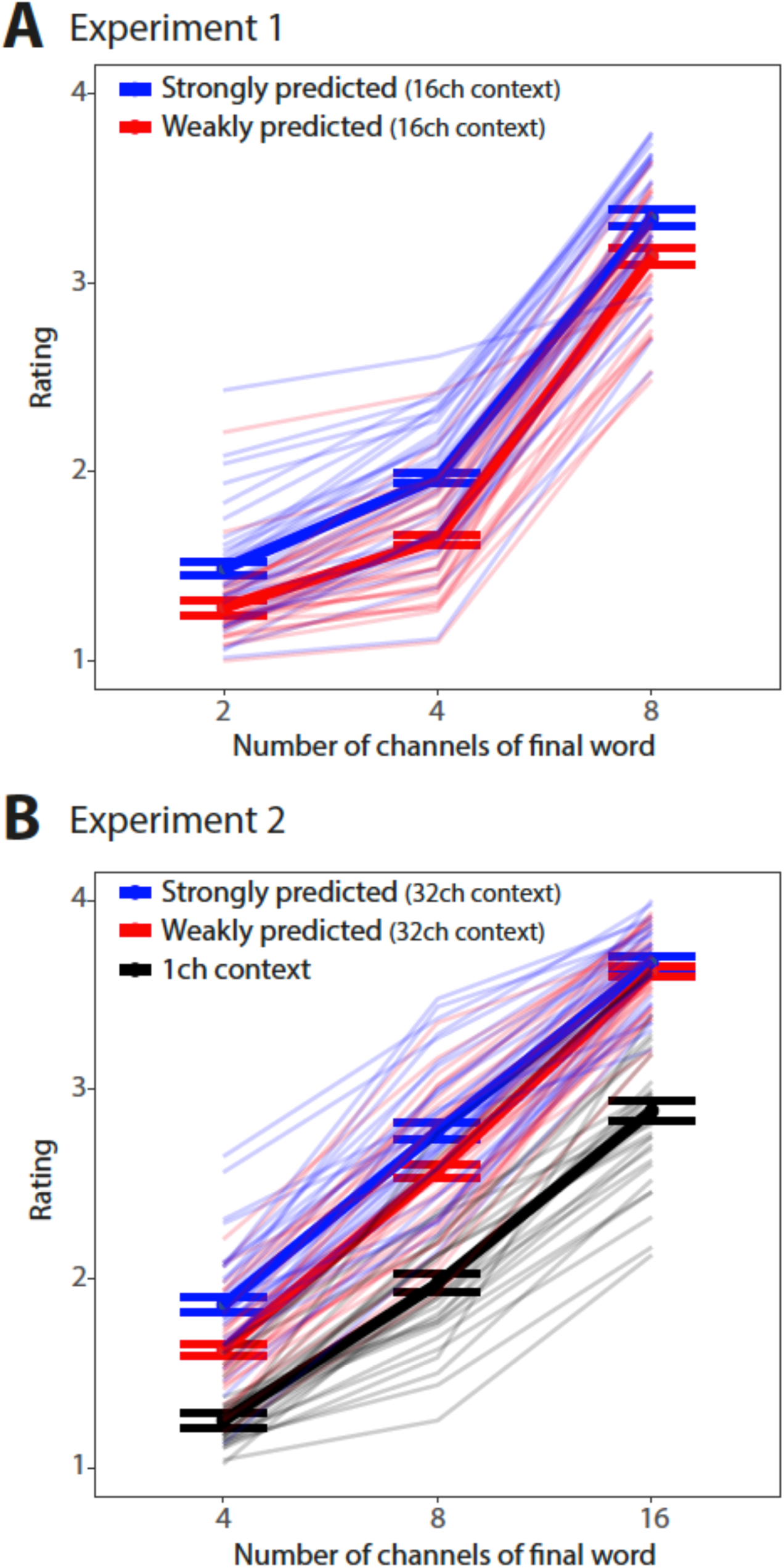
Behavioural results. **A)** Ratings of final word clarity in Experiment 1, with each participant represented by a different line and group averages shown as thicker lines. Error bars show within-subject standard error of the mean [29]. **B)** Ratings of final word clarity in Experiment 2.

#### Cortical tracking: Model accuracies

Previous results show that cortical tracking of speech is driven more by spectral and temporal modulations than other speech features [25]. Moreover, cortical tracking of speech modulations is affected by prediction congruency and signal quality [25]. To confirm that this modulation feature space is also a good model for the present EEG data, we compared model accuracy for the modulation feature space with that of a baseline feature space (broadband envelope) that has been widely used in previous cortical tracking studies [27,30].

On a per-participant basis, model accuracies were averaged over conditions and the 20 sensors with the highest model accuracy were selected. This was done separately for the modulation and envelope feature spaces. The modulation feature space explained EEG neural responses more accurately than the envelope, both for the sentence context period (Figure 5A; t(29) = 13.00, p < .001) and for the final word period (Figure 5B; t(29) = 9.74, p < 001). This confirms that the modulation space is a good model for the present EEG data and therefore in subsequent analyses, we report cortical tracking results using only the modulation feature space.

**Figure 5.**
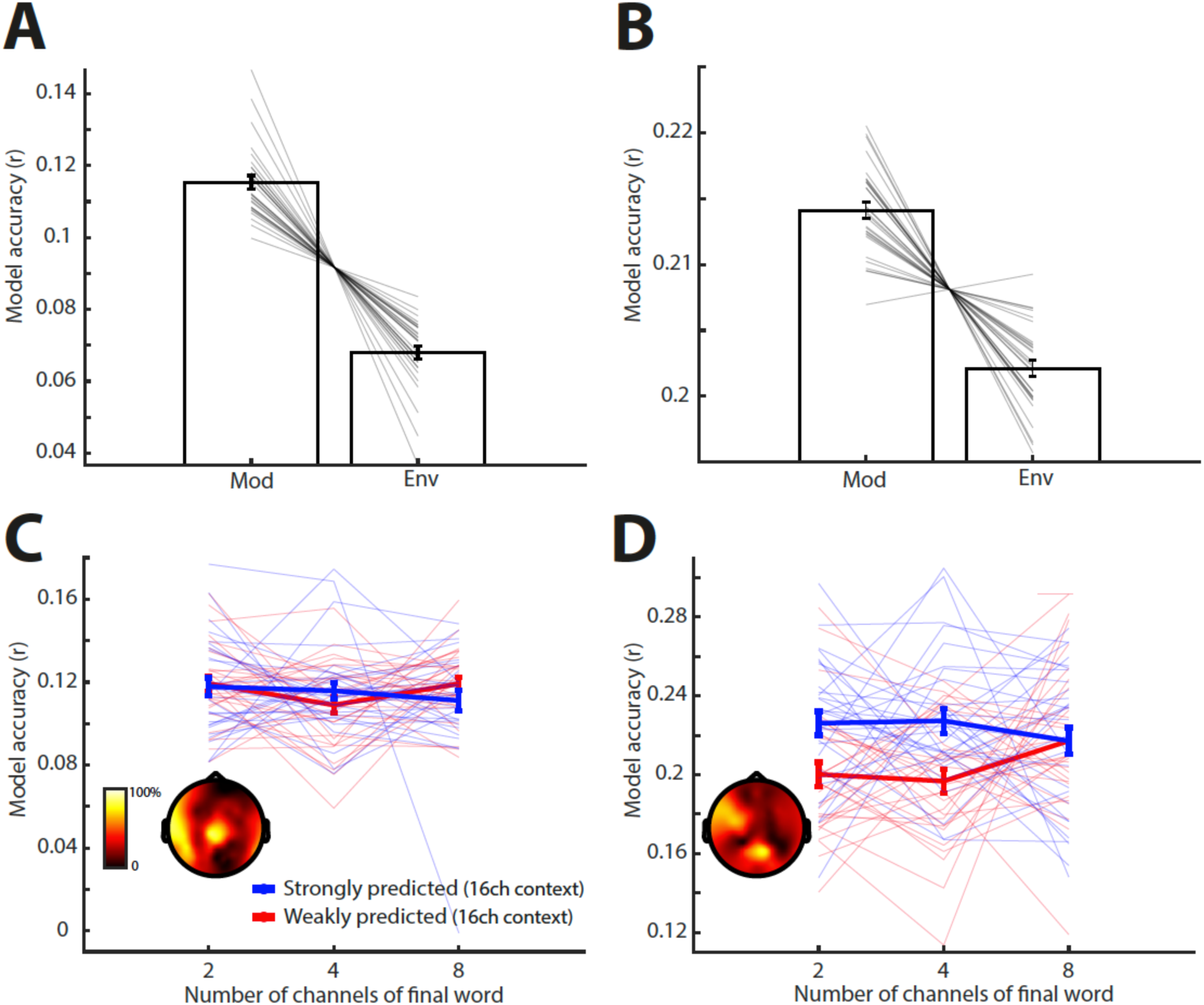
Cortical tracking (model accuracy) results for Experiment 1. **A)** Condition-averaged model accuracies for the modulation and envelope feature spaces, during the sentence context period. **B)** Same as panel A but for the final word period. **C)** By-condition model accuracies for the modulation feature space, during the sentence context period. Sensor selections are indicated in the topography as the percentage of participants for which a sensor was selected. Individual participant data and error bars are shown after removing between-subject variance, suitable for repeated-measures comparisons [29]. Error bars represent standard error of the mean. **D)** Same as panel C but for the final word period.

We next examined how cortical tracking of speech modulations was affected by final word prediction strength and signal quality, focussing first on the sentence context period (Figure 5C). Note that speech features (modulations) were obtained from the original clear versions of the spoken stimuli i.e. before noise-vocoding. This facilitated comparison of model accuracies under different signal quality (vocoder channel) conditions, since any model accuracy differences can be attributed to neural encoding rather than acoustic differences attributable to the signal quality manipulation.

There was no main effect of final word prediction strength (F(1, 29) = .002, p = .96) on model accuracies. Nor was there a main effect of final word signal quality (F(1.98, 57.39) = 1.69, p = .19) or two-way interaction (F(1.96, 56.88) = .96, p = .39). Non-significant effects involving final word signal quality were as expected in Experiment 1 because during sentence context, the signal quality was always the same (16 channels). While in principle there might have been an effect of prediction strength during this period driven by prediction entropy (as explained in Methods), the results in Experiment 1 indicate no impact of prediction on cortical tracking before the onset of the final word.

For the critical final word period (Figure 5D), there was a significant main effect of prediction strength (F(1, 29) = 8.18, p = .008), reflecting larger model accuracies in the strongly predicted versus weakly predicted conditions. There was also a significant interaction between final word prediction strength and signal quality (F(1.99, 57.79) = 3.32, p = .04). From visual inspection of Figure 5D, this significant interaction appears driven by an increase in model accuracy with final word signal quality in the weakly predicted condition, which was not apparent or perhaps even reversed in the strongly predicted condition. However, follow-up testing shows that there was only a trend for a significant signal quality effect in the weakly predicted condition (F(1.64, 47.60) = 2.56, p = .1). In the strongly predicted condition, there was also no significant effect of final word signal quality (F(1.79, 52.04) = .46, p = .62).

To summarise, our cortical tracking analysis for the critical final word period suggests an interaction between prediction strength and signal quality. As explained in the Introduction, an interaction is diagnostic of the prediction error account. However, the precise form of the interaction revealed in the present experiment seems less conclusive. In particular, we do not observe a decrease in speech representation with increasing signal quality when predictions are strong. This is the key pattern that would favour prediction errors over the sharpened signals account (see Figure 2C and 2D).

### Experiment 2

In Experiment 1, we observed robust effects of final word prediction strength and signal quality on neural encoding of speech modulations. However, these results did not clearly distinguish sharpened signal from prediction error accounts. One explanation for this is the range of speech signal quality that listeners heard: the critical final word in each noise-vocoded sentence was presented with 2, 4 or 8 spectral channels. The clearest 8 channel condition is less clear than the equivalent condition in previous work (12 vocoder channels [16,25]) and may have limited the impact of signal quality on neural representations.

Therefore, in Experiment 2 we presented sentences in which the final word was noise-vocoded with 4, 8 or 16 spectral channels, enabling a more extreme manipulation of signal quality (Figure 1B). In addition, we presented the context portion of the sentences with either 32 spectral channels (highly intelligible) or 1 channel (unintelligible). By including a condition in which the sentence context was unintelligible, we can observe neural responses when listeners are unable to make any predictions before the final word is heard (Figure 1B). Consequently, in Experiment 2 the manipulation of prediction strength was also more extreme than in Experiment 1.

#### Behaviour

As shown in Figure 4B, final word clarity ratings were enhanced not only by final word signal quality (F(2, 60) = 1,050.71, p < 0.001) but also by final word prediction strength (F(1.16, 34.69) = 178.45, p < 0.001). There was also an interaction between final word prediction strength and signal quality (F(3.12, 93.68) = 10.21, p < 0.001). These results are essentially identical to that observed in Experiment 1. Follow-up comparisons show that final word clarity ratings in the weakly predicted condition were higher than in the unintelligible context condition (F(1, 30) = 155.36, p < 0.001). In turn, clarity ratings were higher in the strongly predicted than in the weakly predicted conditions (F(1, 30) = 205.03, p < 0.001). This suggests that prediction strength had a graded effect on speech clarity.

#### Cortical tracking: Model accuracies

As in Experiment 1, we examined how prediction strength and final word signal quality affected cortical tracking of speech modulations. This was done separately for sentence context and for final word periods.

For the sentence context period (Figure 6A), there was no main effect of final word signal quality (F(1.97, 59.03) = .59, p = .56) on model accuracies. Nor was there a two-way interaction (F(3.35, 100.65) = .52, p = .69). As in Experiment 1, non-significant effects involving final word signal quality were as expected during the sentence context period i.e. before the final word was heard. However, there was a significant main effect of final word prediction strength (F(1.73, 52.00) = 8.78, p = .001). Visual inspection of the data in Figure 6A indicates that this main effect is caused by higher model accuracies in the weakly predicted and strongly predicted conditions, as compared with the unintelligible context condition (in which no predictions for the final word could be made). This is to be expected because in the unintelligible context condition, the speech was vocoded using 1 channel, compared to 32 channels in the weakly and strongly predicted conditions. Therefore during sentence context, greater signal quality (and, hence, greater intelligibility) results in increased cortical tracking of speech modulations, consistent with previous work [31–33]. Analysis of spectral coherence between the encoding model’s predictions and the observed EEG data (see Methods) may indicate an origin in the low frequency (< 2 Hz) delta band but these coherence effects do not survive FDR correction for multiple comparisons across frequencies (Figure 6C).

**Figure 6.**
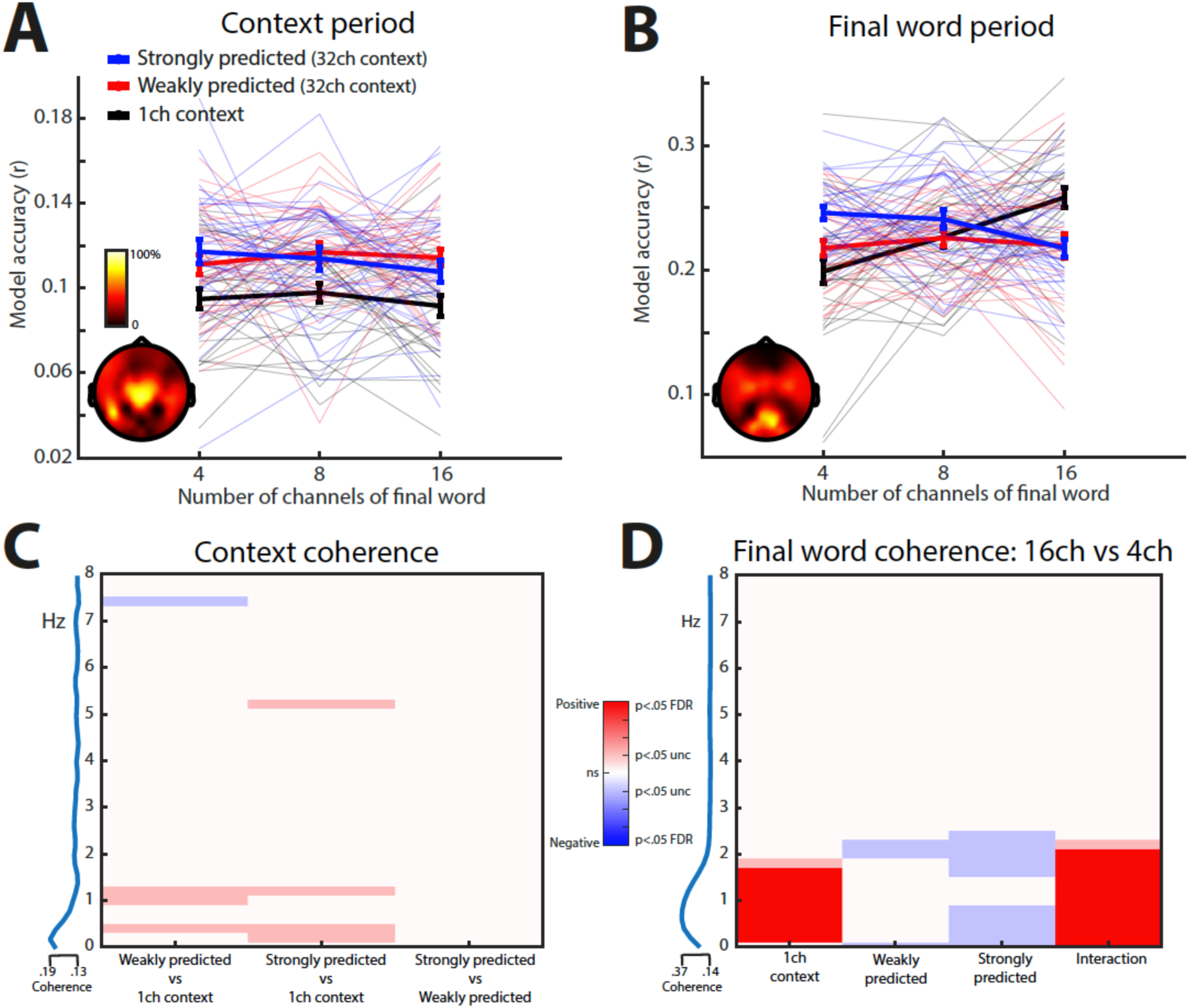
Cortical tracking (model accuracy) results for Experiment 2. **A)** By-condition model accuracies for the modulation feature space, during the sentence context period. Sensor selections are indicated in the topography as the percentage of participants for which a sensor was selected. Individual participant data and error bars are shown after removing between-subject variance, suitable for repeated-measures comparisons (28). Error bars represent standard error of the mean. **B)** Same as panel A but for the final word period. **C)** Model coherence during the sentence context period. The heatmap indicates which frequencies show coherence differences for selected contrasts (positive differences shown in red; negative differences in blue; dark colours represent effects FDR corrected across frequency while light/transparent colours indicate uncorrected differences). The blue line to the left shows model coherence averaged over conditions. **D)** Same as panel C but for the final word period.

For the critical final word period (Figure 6B), there was a significant interaction effect between final word prediction strength and signal quality (F(3.63, 108.93) = 11.54, p < .001). Given this interaction, we performed one-way ANOVAs to examine the effect of final word signal quality separately for 1 channel (unintelligible), weakly predicted and strongly predicted conditions. For the unintelligible context condition, model accuracies increased with increasing final word signal quality (F(1.93, 57.94) = 15.04, p < .001). For the weakly predicted condition, there was no significant effect (F(1.75, 52.47) = .28, p = .73). Critically, in the strongly predicted condition, increasing final word signal quality had the opposite effect on model accuracies as compared with the unintelligible context condition i.e. reduced model accuracies with increasing signal quality (F(1.85, 55.41) = 5.43, p = .008). These results are more consistent with prediction errors than sharpened signals (compare with Figure 2C and 2D). Analysis of spectral coherence (Figure 6D) indicates that this interactive influence of final word prediction strength and signal quality on cortical tracking arises in the low frequency (< 2 Hz) delta band (p < .05 FDR corrected for multiple comparisons across frequencies).

#### Cortical tracking: Model weight analysis

The analyses discussed in the section above reflect cortical tracking of speech modulations over a range of time lags (0 to 450 ms). To examine the timecourse of the neural interaction between final word prediction strength and signal quality, we analysed the encoding model weights i.e. temporal response functions (TRFs). We first analysed TRFs in sensor space, summarised as the RMS amplitude over sensors. Sensor selections were the same as those used for the model accuracy analysis (previously shown in Figure 6A and 6B). All effects are reported after applying a p < .05 threshold with FDR correction for multiple comparisons over lag and modulation features.

As shown in Figure 7A, TRF weight amplitudes during the sentence context period were larger in the weakly and strongly predicted conditions versus the unintelligible (1 channel) context condition. This effect closely followed stimulus change (-50 to 50 ms lag; shown as red patches) and extended into negative latencies. This latter observation may suggest a predictive neural signal, reflecting e.g. pre-activation of speech features [34]. However, as noted by Teoh et al. [35], encoding models may not always be as highly temporally resolved as they may appear, due to the autocorrelational structure of speech. We therefore refrain from drawing strong conclusions concerning negative latency effects. Regardless, the increase in TRF weights for the weakly and strongly predicted conditions (in which speech was noise-vocoded using 32 channels), suggests that differential tracking of speech due to spectral detail occurs within 50 ms of stimulus change. This finding is consistent with previous work showing early cortical tracking of acoustic features (e.g. speech envelope tracking within 30 ms [36]). At a later latency (approximately 175 ms), TRF amplitudes were modulated in the opposite direction i.e. decreased response for weakly/strongly predicted versus unintelligible context conditions (shown as blue patches).

**Figure 7.**
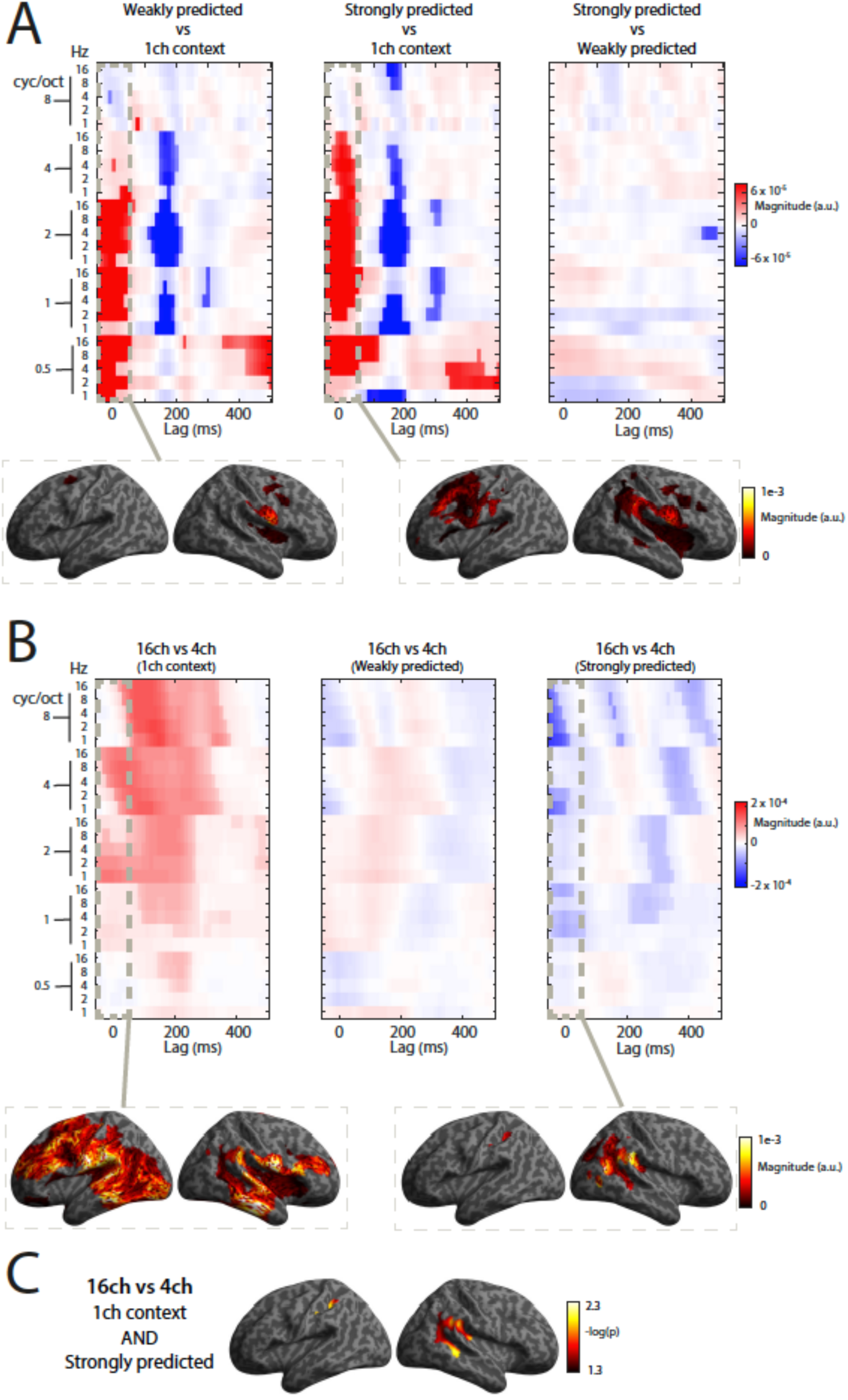
TRF model weights in Experiment 2. **A)** TRF model weights during the sentence context period. The heatmap indicates which speech features and lags show weight differences for selected contrasts (positive differences shown in red; negative differences in blue; dark colours represent effects FDR corrected across speech features while light/transparent colours indicate uncorrected differences). Brain plots overlay source-transformed weights onto an MNI-template cortical surface, averaged from -50 to 50 ms lags and over selected speech features (2 to 8 cycles per octave and over all temporal modulations). **B)** Same as panel A but for the final word period. **C)** Same as panel B but for the conjunction contrast [ 16ch > 4ch (1ch context) ] AND [16ch < 4ch (Strongly predicted) ].

As shown in Figure 7B, TRF weight amplitudes during the final word period show a pattern highly consistent with the previous model accuracy analysis. For the unintelligible context condition, TRFs increased with increasing final word signal quality while the opposite occurred in the strongly predicted condition i.e. reduced TRFs with increasing signal quality. As with the previous model accuracy analysis, there was no effect of signal quality in the weakly predicted condition. These results, as with the model accuracy analysis in Figure 6B, are more consistent with the prediction error than sharpened signal account. Effects of signal quality on TRFs are first apparent around stimulus change (-50 to 50 ms lag) but continue into later lags (until around 400 ms).

We further localised the TRFs to the underlying brain sources (see Methods). For this source analysis, we first averaged the weights over speech modulation features, between 2 and 8 cycles per octave. It is this spectral modulation range that is most impacted by noise-vocoding (degrading spectral fine structure [25]) and therefore might signal mismatch with top-down predictions. This modulation range also shows an interaction between final word prediction strength and signal quality in sensor space (shown previously in Figure 7B). We further focus on the earliest period (-50 to 50 ms) as this period may better capture the primary consequences of our experimental manipulations (as opposed to secondary post-perceptual effects). For both the sentence context and final word periods, changes in TRFs localise to perisylvian cortex bilaterally including the superior temporal gyrus (shown in the cortex renderings at the bottom of Figure 7A and 7B). This region has previously been implicated as a source of prediction errors during single word processing [5,16,37].

To more selectively isolate the interaction effect between prediction strength and signal quality during the final word period, we tested for the conjunction of the signal quality effect (16 versus 4 channels) in the unintelligible and strongly predicted conditions. This revealed a more focal effect in the posterior portion of the right superior temporal gyrus as well as the left inferior partial lobule (Figure 7C).

### Computational simulations: Timecourse analysis

As discussed above, the EEG results of Experiment 2 favour a prediction error account. In particular, the impact of final word signal quality when speech is completely unpredicted (in the unintelligible context condition) and when speech is strongly predicted, is highly convergent in the simulated and observed data. However, a discrepancy between the simulated and observed data occurs in the weakly predicted condition. Simulated prediction errors at word onset increase with signal quality in the weakly predicted condition. However, cortical tracking does not vary (compare Figure 2D and Figure 6B). One possible explanation for this is apparent in the model outputs depicted in Figure 2A for the final word in “In sailing to have to handle a lot of **water**”. For this item, predictions for the final word are weak (prior word probability of .03). However, by segment position 2 (after hearing the vowel /O:/), the unfolding sensory evidence is sufficient to rule out alternative word candidates and provide support in favour of the stimulus word (i.e. a higher posterior probability for the word *water*). This in turn strengthens predictions for subsequent stimulus segments in positions 3-4. Therefore, although this word is weakly predicted at word onset, it becomes more strongly predicted as the stimulus unfolds in time. This temporal build-up of prediction strength (in what is nominally a weakly predicted condition) could in turn produce a prediction error response similar to that observed in the strongly predicted condition. When averaged over segment position, changes in prediction error attributable to signal quality would then appear as an intermediate pattern between that of the unintelligible context condition (in which no predictions are possible) and that of the strongly predicted condition. This pattern could also be reflected in the EEG TRF analysis, which measures the time-averaged relationship between speech and neural responses i.e. over all segment positions.

To test this possibility, we examined simulated prediction errors not just at word onset (as previously depicted in Figure 2D), but for multiple segment positions (shown in Figure 8A). For completeness, we did the same for the sharpened signal simulations (Figure 8B). Note that the temporal dimension for this analysis is fundamentally different to the temporal (by lag) dimension of the TRF analysis (previously depicted in Figure 7). The by-lag analysis of TRF weights examined the magnitude of neural tracking at different time delays between speech and neural responses, averaged over the temporal extent of the final word (as explained above). By contrast, the present simulations examine processing segment-by-segment within the final word.

**Figure 8.**
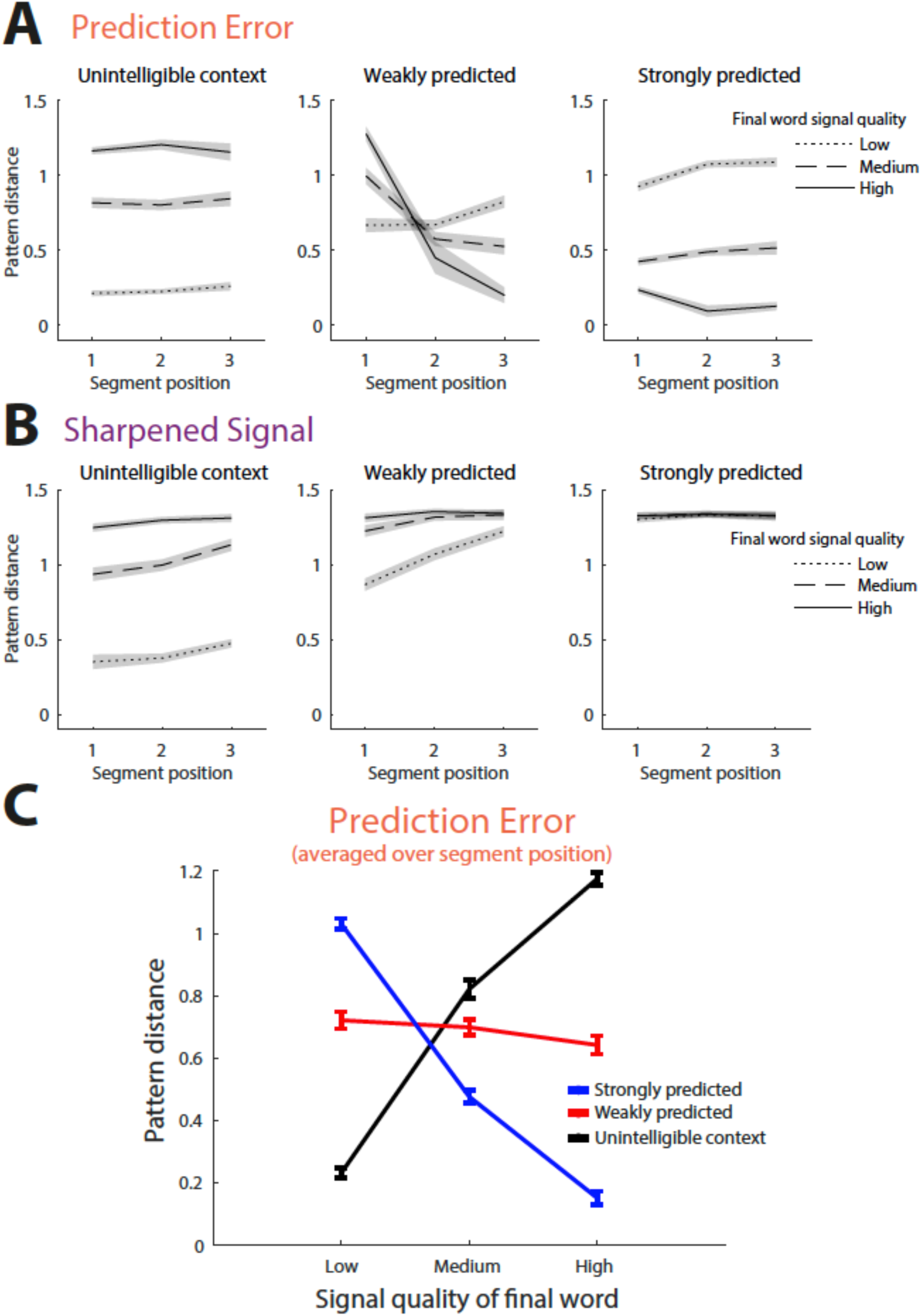
Computational simulations by stimulus segment position. **A)** Mean pattern distances for simulated prediction errors as a function of stimulus segment position. Error bars show the standard deviation. **B)** Same as panel A but for sharpened signals. **C)** Pattern distances for simulated prediction errors averaged over segment position.

Consistent with the explanation above, prediction errors in the weakly predicted condition decrease rapidly during segment positions 2 and 3 when final word signal quality is high (16 channels). This decrease comes about because the increased sensory evidence in the high signal quality condition enables more accurate predictions as the stimulus word builds up over time. This produces a response in the weakly predicted condition that more closely resembles the strongly predicted condition. When averaged over segment position (shown in Figure 8C), prediction errors in the weakly predicted condition remain unchanged with increasing signal quality, as observed in the EEG data (compare Figure 8C with Figure 6B).

## Discussion

In this study we sought to determine how top-down predictions are combined with the sensory signal during perception of connected speech. In two experiments, we observed additive effects of final word signal quality and prediction strength on clarity ratings. Yet these manipulations had an interactive influence on final word neural tracking, which was most clearly observed in Experiment 2. This interactive influence on neural tracking emerged rapidly (within 50 ms of the ongoing speech signal), in low frequency delta frequencies and localized to perisylvian cortex bilaterally, including the superior temporal gyrus.

We tested whether neural responses reflect sharpened signals or prediction errors. As with previous work [16,25], these computations were instantiated through model simulation, which confirmed distinct neural outcomes for sharpened signals versus prediction errors. In Experiment 1, while final word neural tracking was reliably modulated by prediction strength, we failed to observe reliable effects of signal quality. Overall, the pattern of results did not clearly favour either computation. However, in Experiment 2 when signal quality and prediction strength varied more extremely, we observed a clear crossover interaction on neural tracking of a form that appears highly consistent with prediction errors. Taken together these results suggest that cortical responses predominantly reflect the difference between neural representations of heard and predicted speech sounds.

The present results complement previous work demonstrating neural signatures of prediction error using fMRI [16] and MEG [25]. However, in this previous work, listeners heard isolated words and made predictions using external written cues. Thus, the present study goes beyond this previous work in demonstrating the operation of prediction error computations in a more ecological setting. The present paradigm is more naturalistic not only because of the use of sentences (versus single words) but also because listeners’ predictions were based on information intrinsic to the speech signal rather than on external (i.e. written) cues. Our findings parallel other recent work demonstrating prediction errors from speech-intrinsic information [37], albeit in the context of single word listening.

Our work joins several previous studies that have also investigated how speech predictability modulates neural tracking of connected speech [38–43]. Many of these studies make use of the “pop-out” effect, which refers to the marked increase in perceptual clarity that listeners experience when degraded speech is presented shortly after hearing the undegraded version or seeing a written transcription [44]. Using this paradigm, previous intracranial [41] and scalp [38–40] neurophysiology studies are generally convergent in suggesting that top-down predictions enhance neural tracking (although see [42]). Aside from the non-ecological nature of this experimental design (as it is rare in natural listening to hear exact repeats of an utterance), it is notable that most of these studies presented speech at a fixed (low to intermediate) level of signal quality. However, when signal quality is low, increased neural tracking of predicted versus unpredicted speech is in fact consistent with both sharpened signal and prediction error accounts (as can be seen in the computational simulations depicted in Figure 2C and 2D). To adjudicate between sharpened signals and prediction errors, we suggest that it is necessary to jointly manipulate top-down predictions and bottom-up signal quality. Furthermore, as the results of Experiment 1 demonstrate, the bottom-up signal quality must be manipulated over a sufficiently varied range, from highly distorted to highly intelligible speech. Based on Experiment 1 alone, one might favour a sharpened signal account, since the observed pattern (enhanced neural representation of strongly versus weakly predicted stimuli) has previously been interpreted as reflecting sharpened signal computations [45]. However, by manipulating signal quality more extremely in Experiment 2 and examining the impact on neural and simulated representations, the present study demonstrates a cross-over interaction between prediction strength and signal quality that clearly favours the prediction error account.

A key feature of our study was the use of time-limited noise-vocoding, in which differing levels of noise-vocoding were applied to the sentence context and final word periods. This was critical to enable a situation in which strong top-down predictions were met with highly distorted speech input, as was the case in the low signal quality, strongly predicted condition. This would not have been possible had we applied low signal quality noise-vocoding to the entire sentence [46,47], since final words in strongly predicted sentences would have become less predicted following a low signal quality (less intelligible) sentence context. Therefore, it is only through time-limited noise-vocoding that we were able to independently manipulate final word prediction strength and signal quality, which as discussed above is critical for testing sharpened signal and prediction error accounts.

In addition to the analysis of model accuracies, for Experiment 2 we also analysed the model weights linking speech features with EEG responses at specific time lags i.e. TRFs [27,48]. The interactive influence of prediction strength and signal quality on the TRFs closely lagged stimulus change (within 50 ms) and localised to acoustic-phonetic regions in the superior temporal lobe. Early latency effects in auditory regions support the proposal that prediction errors are evident already at lower sensory levels of processing [49–51] and may signal graded (acoustic) mismatch between expected and heard speech sounds [25,52].

Complementing the temporal analysis discussed above, we also examined final word neural tracking in individual neural frequencies. We observed neural tracking in low frequency delta activity which peaked at an ‘ultra low’ peak frequency of 0.6 Hz. Within this low frequency delta range, we observed the prediction error interaction between final word signal quality and prediction strength. This low frequency locus is notable as many neural tracking studies highpass filter neural responses above 1 Hz, which may attenuate a 1 Hz prediction error signal given filter transition bandwidths [53]. One of the notable exceptions to this is the MEG study reported by Donhauser and Baillet [20] which demonstrated a low frequency delta signal that tracked speech predictability (phoneme surprisal) in TED talks (see also [23,24,54,55]). Considered in conjunction with the present results, we suggest that low frequency neural activity may be particularly important for signalling prediction error.

Finally, our computational simulations plausibly explain why neural tracking in the weakly predicted condition diverges from the pattern we had initially expected under the prediction error account. When examining simulated prediction errors at final word onset, increasing signal quality should result in elevated prediction errors in the weakly predicted condition, much like in the unintelligible context condition when no prediction is possible. However, in both experiments we failed to observe a reliable effect of signal quality on final word neural tracking in this condition. Follow-up simulations examined prediction error not only at word onset but also at later segments. In the weakly predicted condition, these simulations produced a pattern that, when averaged across segment positions, more closely resembled the observed EEG data. This occurs because, although the final word may be weakly predicted at its onset, it becomes more strongly predicted as sensory evidence accumulates, simultaneously reducing segment predictions for competitor words.

These temporal dynamics mimic the lexical competition behaviour of cognitive models of spoken word recognition [56–58]. They are also reminiscent of the notion of ‘explaining away’ in predictive coding models of vision [8,59]. In these models, new visual input (e.g. a face or written word) results in an iterative minimisation of prediction errors by top-down predictions until competing interpretations are suppressed. Notably however, for a dynamic signal such as speech, the cortex must not only minimise prediction errors for what has just been heard but also for subsequent word segments [60]. It is this latter aspect that is revealed by our simulations – prediction errors at word onset leading to updated predictions for future word segments.

One limitation of the encoding analysis method utilised here is that we are unable to examine neural tracking segment-by-segment as this would result in too few observations for the linear regression. This precludes us from directly testing whether the computational simulations match the EEG data on a segment-by-segment basis. Future work is therefore needed to enable the linking of neural responses with specific moments in the speech signal, potentially using alternative methods such as representational similarity analysis [61]. The temporal dynamics of the present simulations suggest that this avenue of research is a potentially powerful means of understanding and testing neural implementations of predictive processing.

## Materials and Methods

### Ethics statement

All participants gave written informed consent before completing the study and were reimbursed for their time. The study was approved by the University of Sussex Science and Technology Cross Schools Research Ethics Committee.

### Participants

All participants were native speakers of British English and had no history of hearing impairment or neurological disease based on self-report. All participants were naive to the purposes of the experiment. Different participants were recruited for the two experiments. Participants were required to be between 18 and 35 years of age to take part. The participants that were tested were between 18 and 29 years of age.

For Experiment 1, 34 participants were tested. One participant was excluded due to failure to understand task instructions, and three participants were excluded due to technical issues with the EEG recording. Therefore 30 participants were included in the final analysis, of which 20 were female (mean = 21.06 years of age, SD = 2.15). Of the participants included in the final analysis, one participant was left-handed.

For Experiment 2, 35 participants were tested. Four participants were excluded due to technical issues with the EEG recording. Therefore 31 participants were included in the final analysis, of which 19 were female (mean = 19.32 years old, SD = 1.05). Of the participants included in the final analysis, three participants were left-handed.

### Spoken stimuli

216 sentences were presented to each participant in spoken format. The sentences were taken from Peelle et al. [62], a study which reported completion norms from at least 100 participants for English written sentences of eight to ten words in length. From these completion norms, Peelle et al. [62] report final word cloze probabilities which are a widely used measure of word predictability [63,64] and are obtained by computing the proportion of participants who completed the sentence with a particular word.

For the ‘strongly predicted’ condition, we selected 108 sentences in which the final word had a high cloze probability (ranging between 0.66 and 0.99; mean = 0.84, SD = 0.10). For example, “He was late because he had gotten a flat **tyre**”, which has a cloze probability of 0.99. For the ‘weakly predicted’ condition, we selected 108 sentences in which the final word had a low cloze probability (ranging between 0.02 and 0.10; mean = 0.07, SD = 0.02). For example, “In sailing you have to handle a lot of **water**”, which has a cloze probability of 0.03 (because participants tend to predict the alternative endings *ropes*, *wind* or *waves*). All sentences were plausible and semantically coherent. Distributions of cloze probabilities are shown in Figure 3A. Further examples of sentences are shown in Figure 1A.

In addition to cloze probabilities, Peelle et al. [62] report a second measure of final word predictability: entropy. Different to cloze probability (which relates to the prediction for a single word), entropy relates to the distribution of predictions over multiple possible word continuations and captures the constraint provided by sentence context prior to the final word (measured in bits). Formally, entropy (*H*) is computed as follows:

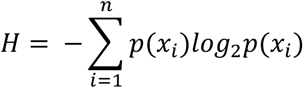

where p(x) is the probability distribution over words (measured here as the proportion of participants completing the sentence with each word continuation).

Items with high entropy indicate sentence contexts that offer weak constraint as to the identity of the final word whereas items with low entropy indicate sentence contexts with strong constraints. In natural speech, these two measures of predictability are negatively correlated [65], although there are cases when both entropy and cloze probability can be low. This occurs when hearing an unexpected word following a constraining (low entropy) context. For example, after hearing the words “The little girl tripped and immediately began to …”, listeners will strongly predict the word *cry* and make only weak predictions for other words. Accordingly, this sentence context is highly constraining, and entropy is low. However, if this sentence ends instead with the word *fall*, the final word is relatively unpredicted (low cloze probability) even though the context is highly constraining. Thus, while both cloze probability and entropy are closely related measures, they are not identical and together provide a more complete characterisation of speech predictability.

In the current stimulus set, log cloze probability and entropy had a Pearson’s correlation coefficient of -.74. In the strongly predicted condition, entropy was relatively low (ranging between 0.08 and 1.62; mean = 0.91, SD = 0.45) but in the weakly predicted condition, entropy varied substantially (ranging between 0.55 and 5.43; mean = 2.85, SD = 1.04; see Figure 3A). So while the final word in the weakly predicted condition always had a low cloze probability, this followed contexts varying in constraint.

After selecting the sentences, we recorded spoken versions from a male speaker of southern British English. Recordings were acquired using an Audio Technica AT2035 condenser microphone and saved as 16-bit, 48 kHz wav files. The duration of the sentences ranged from 2207 to 3978 ms (mean = 3006, SD = 376).

As shown in Figure 3B, the acoustic properties of the final word were well matched between strongly and weakly predicted conditions. Using independent samples t-tests to compare the strongly and weakly predicted conditions, we confirmed that there were no reliable differences in final word duration (t(214) = 1.76, p = .080), intensity (t(214) = 1.09, p = .276) and fundamental frequency (t(214) = .228, p = .820).

We used noise-vocoding to vary signal quality. The noise-vocoding procedure [66] superimposes the temporal-envelope from separate frequency regions in the speech signal onto white noise filtered into corresponding frequency regions. This allows parametric variation of signal quality (spectral detail), with increasing numbers of channels associated with increasing intelligibility. Vocoding was performed using a custom Matlab script (The MathWorks Inc), with the spectral channels spaced between 70 and 5000 Hz according to Greenwood’s function [67]. Envelope signals in each channel were extracted using half-wave rectification and smoothing with a second-order low-pass filter with a cut-off frequency of 30 Hz. The overall RMS amplitude was adjusted to be the same across all audio files.

We adapted the standard noise-vocoding procedure to manipulate final word signal quality independently of sentence context (i.e. the portion of the signal leading up to final word onset). In Experiment 1, the sentence context portion of the sentence was noise-vocoded using 16 spectral channels and therefore was highly intelligible (see Figure 1A).

The signal quality of the final word was manipulated by noise-vocoding using 2, 4 or 8 spectral channels. In Experiment 2, sentence context was noise-vocoded using either 32 channels (fully intelligible) or 1 channel (unintelligible) whilst the final word was noise-vocoded using 4, 8 or 16 channels (see Figure 1B).

To achieve this separate noise-vocoding of the context and final word periods of the sentence, we performed the following procedures. First, we noise-vocoded the entire sentence (e.g. using 16 channels). Second, we noise-vocoded the same sentence using a different number of channels (e.g. 2 channels). Third, we mixed the two signals using a Hann window of 10 ms to transition from the 16 channel to 2 channel versions. The combination of a Hann window and the stochastic nature of the noise carrier ensured that there was no perceptible discontinuity between the two signals.

In Experiment 1, the assignment of sentences to final word signal quality conditions was counterbalanced across participants. In Experiment 2, the assignment of sentences to 32 channel versus 1 channel context conditions was also randomised across participants. For the 1 channel context condition, sentences were drawn equally from the strongly and weakly predicted conditions.

### Procedure

Participants completed a clarity rating task in which they were required to rate the clarity of the final word in each sentence. Participants were instructed to “*decide how clear the final word is on a scale of 1 to 4. This is a comparative task, so 1 = least clear when compared to the other words; 4 = most clear when compared to the other words.*” For Experiment 1, manipulations of final word signal quality (2 / 4 / 8 channels) and final word prediction strength (strongly / weakly predicted) resulted in a 3 × 2 factorial design. For Experiment 2, manipulations of final word signal quality (4 / 8 / 16 channels) and final word prediction strength (strongly predicted / weakly predicted / 1 channel context) resulted in a 3 × 3 factorial design.

During trials, a fixation cross was always shown on screen, which consisted of a white cross on a grey background. Each trial commenced with the presentation of a spoken sentence, followed 1000 ms later with a cue for the participant to respond (consisting of the fixation cross changing colour from white to black). Responses were recorded from the participants left hand using the keyboard. The experimenter checked before the experiment began that the participant was able to enter their intended response without looking at the keyboard. Subsequent trials began 2000 (±0–100) ms after participants responded.

Both Experiments 1 and 2 consisted of six blocks of 72 trials each. In blocks 1 - 3, all 216 unique sentences were heard once in randomised order, with each block containing an equal number of trials per condition. In blocks 4 - 6, each stimulus was repeated but in a different randomised order. Before the experiment, participants completed two practice sessions, each consisting of ten sentences. These contained equal numbers of strongly predicted and weakly predicted trials.

### Data acquisition and preprocessing

Sounds were presented via Sennheiser HD 380 Pro headphones and an M-Audio M-Track solo external sound card. Stimulus delivery was controlled using PsychoPy software (v2022.1.1 Open Science Tools Ltd).

EEG data were recorded with ASA-Lab using a 64 channel ANT Neuro amplifier with a sampling rate of 1000 Hz and a 64 channel Waveguard EEG cap (ANT Neuro, Enschede), with electrodes placed according to the extended 10-20 system and offline referenced to the average across electrodes. Pre-processing of the EEG data was performed using the SPM12 toolbox (Wellcome Trust Centre for Neuroimaging, London, UK) in MATLAB R2020b (Mathworks, Inc. Natick, MA, USA) and data analyses were performed using custom MATLAB scripts. EEG data were downsampled to 250 Hz and noisy channels reconstructed from neighbouring channels. DC offsets were removed by subtracting the mean from each channel. Robust detrending was then used to remove slow drifts [68]. Signal components corresponding to line noise (50 Hz) and the monitor refresh rate (85 Hz) were removed using Zapline [69].

Independent components analysis was used to remove activity related to blinks and eye movements. Identification of eye artefact components was achieved using an automatic procedure based on multiple linear regression. Specifically, the signal from the vertical and horizontal EOG sensors were used as predictor variables to regress against each ICA component time-series. Prior to regression, EOG and ICA signals were first z-scored such that each signal had a mean of zero and standard deviation of one. ICA components for which R2 (variance explained) was above 0.25 was identified as eye-related activity and subsequently projected out from the data.

For some participants, we removed blocks where the recording was excessively noisy. For Experiment 1, a total of five blocks were removed (two blocks for two participants and one block for one participant). For Experiment 2, one block was removed for each of two participants.

### Encoding Models

As in Sohoglu and Davis [25], we used ridge regression to measure cortical tracking of speech features [36,48,70–72]. Our analysis approach is near-identical to the one used in Sohoglu and Davis [25]. The analysis described below was done separately for the context period (sentence onset to final word onset) and final word period (final word onset to sentence offset).

Before model fitting, the EEG data were epoched at matching times as the feature spaces (see below), and outlier trials removed. We then used the mTRF toolbox [27] to fit an encoding model that mapped from the stimulus features to the neural time-series observed at each EEG sensor and at multiple time lags:

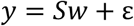

Where y is the neural time-series recorded at each sensor, S is an N_samples_ by N_features_ matrix defining the stimulus feature space (concatenated over different lags), w is a vector of model weights and ε is the model error. Model weights for positive and negative lags capture the relationship between the speech feature and the neural response at later and earlier timepoints, respectively. Encoding models were fitted with lags from -100 to 550 ms and tested with a more restricted range of lags from 0 to 450 ms, to avoid contributions from possible edge artefacts that can affect model weights.

To control for overfitting, we used ridge regression and varied the lambda parameter (over 21 values as follows: 2^0^, 2^1^, 2^3^, … 2^20^) which controls the degree to which strongly positive or negative weights are penalised [27]. This lambda parameter was optimized using a leave-one-trial-out cross-validation procedure. For each feature space, sensor, condition, and participant, we fitted models using data from all but one trial. We then averaged the model weights across trials and used the result to predict the EEG response of the left-out trial. We computed model accuracy as the Pearson correlation between the predicted and observed data and repeated this procedure such that model accuracies were obtained for all trials. Before optimizing lambda, model accuracies were averaged across trials and across sensors. The optimal lambda value was then selected as the mode of the model accuracy distribution over participants and conditions [41]. Two feature spaces were tested:

1. Spectral and temporal modulations, capturing regular fluctuations in energy across the frequency and time axes of the spectrogram [28,73]. We used the NSL toolbox in Matlab (http://nsl.isr.umd.edu/downloads.html) to first compute a 128 channel ‘auditory’ spectrogram (with constant Q and logarithmic centre frequencies between 180 and 7040 Hz), before filtering with 2D wavelet filters tuned to spectral modulations of 0.5, 1, 2, 4, and 8 cycles per octave and temporal modulations of 1, 2, 4, 8, and 16 Hz. All other parameters were set to the default (frame length = 12.5 ms, time constant = 12.5 ms and no nonlinear compression). Note that temporal modulations can be positive or negative, reflecting the direction of frequency sweeps (i.e. upward versus downward [74]). This default representation is a very high-dimensional feature space: frequency × spectral modulation × temporal modulation × temporal modulation direction with 128 × 5 × 5 × 2 = 6400 dimensions, each represented for every 12.5 ms time sample in the speech file. We therefore averaged over the 128 frequency channels and positive- and negative-going temporal modulation directions. The resulting feature space was a time-varying representation of spectral and temporal modulation content comprised of 25 features (five spectral modulations x five temporal modulations). The method and parameters for extracting this feature space are identical to Sohoglu and Davis [25].
2. Broadband envelope, capturing time-varying acoustic energy in a broad 180– 7040 Hz frequency range. This feature space was obtained by summing the spectral channels in the auditory spectrogram described above.

These feature spaces were obtained from the original clear versions of the spoken stimuli (i.e. before noise-vocoding) and downsampled to 80 Hz to match the EEG sampling rate. The feature spaces were then z-score transformed such that each time-series on every trial had a mean of zero and standard deviation of 1.

To examine cortical tracking in individual neural frequencies, we computed the coherence between the encoding model predictions and the observed EEG data (see [20]), using the same cross-validation scheme as for the model accuracy analysis. Spectral coefficients were estimated using the Fourier transform and a Hanning taper over a 5 second window. We analysed model coherences at individual neural frequencies in steps of 0.2 Hz within the delta/theta range (0-8 Hz), which previous work has implicated in cortical tracking of speech features [20,40].

### Source reconstruction of encoding model weights

To determine the underlying brain sources of the encoding model effects, we used a distributed method of source reconstruction, implemented within the parametric empirical Bayes framework of SPM12 [75–77]. The forward model was computed using a Boundary Element model and sensor positions projected onto an MNI space template brain (sensor positions were as specified by the ANT neuro template). For inversion of the forward model, we used the ‘LOR’ routine in SPM12, which assumes that all sources are activated with equal *a priori* probability and with weak correlation to neighbouring sources. This was applied to the encoding model weights from -50 to 500 ms lags, averaged over all temporal modulations and over 2 to 8 cycles per octave.

Source solutions were constrained to be consistent across subjects, which has been shown to improve group-level statistical power [76]. In brief, this procedure involves 1) realigning and concatenating sensor-level data across subjects 2) estimating a single source solution for all subjects 3) using the resulting group solution as a Bayesian prior on individual subject inversions. Thus, this method exploits the availability of repeated measurements (from different subjects) to constrain source reconstruction. Importantly, however, this procedure does not bias activation differences between conditions in a given source.

To analyse condition-wise effects on source activity, we focus on the earliest period (-50 to 50 ms) when an interaction between prediction strength and signal quality was observed in sensor space. Source power from 0 to 40 Hz within this period was summarised using a Morlet wavelet projector [78]. Statistical maps of source activity are displayed using an FDR corrected threshold of p < .05.

### Computational modelling

To guide interpretation of the neural data, we simulated sharpened signal and prediction error representations using a Bayesian model of spoken word recognition [79], as implemented in Sohoglu et al. [37]. At the word level (illustrated as the green word clouds in Figure 2A), we computed a posterior distribution over a lexicon of 73,120 words, by combining word priors and likelihoods using Bayes rule. Word priors were based on the cloze response probabilities provided by Peelle et al. [62] while word likelihoods were computed by combining segment input probabilities (derived from acoustic similarities; see below for further details). This was done iteratively at each segment position in the stimulus word.

Consider for example the sentence context “In sailing you have to handle a lot of …”. While *water* (SAMPA transcription: / w O: t @ /) is a plausible completion of this sentence, this word is only weakly predicted (p = 0.03) in comparison with other words such as *ropes* (p = 0.32), *wind* (p = 0.27) or *waves* (p = 0.12). Following /w/ at segment position 1 in the stimulus, word likelihoods (i.e. the sensory evidence) are updated. When combined with word priors, this leads to higher posterior probabilities for words starting with the segment /w/ (e.g. *water*, *wind*, *waves* etc.) and lower probabilities for other words (e.g. *ropes*, *sails* etc.). At this point, *water* is not yet the most probable word because *wind* also matches the sensory input and has a stronger prior probability (p = 0.27 versus p = 0.03). However, following a further phoneme at segment position 2 (the vowel /O:/), word likelihoods are again updated and combined with word priors, making *water* the most probable word. In this way, we computed a word posterior distribution at each segment position in the stimulus word.

The above computations perform Bayesian inference at the word level. In predictive processing models, top-down predictions from higher hierarchical levels are propagated to lower levels [6,10,80]. Indeed, previous evidence suggest that cortical responses to speech are well explained by predictive computations occurring at the segment level [5,81]. To compute predictions for each upcoming segment (shown as green phonetic symbols in Figure 2A), we summed lexical level posterior probabilities over words sharing the next segment. This enabled the expression of predictions for each of 44 phonemes (i.e. the model’s phonological inventory) at each segment position. Returning to the example above, following /w/ at segment position 1, predictions are made for the segment /I/ (for *wind*) but also /O:/ (for *water*). Because *wind* is more strongly predicted by the sentence context than *water* (p = 0.27 vs p = 0.03), the next segment prediction for /I/ is also stronger than for /O:/. By segment position 2 however (after the vowel /O:/), sufficient sensory evidence has accumulated to favour *water* as the most probable word candidate. In turn, this leads to a strong prediction that the next segment is /t/. This example illustrates that predictions for speech segments in the model are determined not only by word priors from sentence context but also by accumulating sensory evidence as the speech signal unfolds.

Following the computation of these segment predictions, we generated two model outputs differing in the way that sensory evidence for each of the 44 phoneme segments in the model’s phonological inventory was combined with the segment predictions. Under the sharpened signals simulation, the model output was calculated by multiplying segment predictions with segment input probabilities (normalising to sum to one). Under the prediction error simulation, the model output was calculated by subtracting segment predictions from the segment input probabilities. These computations were performed iteratively for each segment position in the unfolding stimulus word.

Therefore, at each segment position in the unfolding stimulus word, sharpened signals or prediction errors comprise a vector of 44 segments (the model’s phonological inventory). To summarise these vectors at each segment position, we computed the Euclidean pattern distance between the vectors for all pairs of items within each condition (see [37]). For example, we computed the distance between the vector for the onset segment of *water* with that of *beagle*; between *water* and *pass*; between *water* and *money*; between *beagle* and *water*; between *beagle* and *pass* etc. Pattern distances were computed for segment positions 1-3 only since fewer stimuli were available for later segment positions. We then averaged the pattern distances for all stimuli presented to each participant.

This pattern distance analysis of our simulations quantifies the amount of speech information content in sharpened signals and prediction errors, with larger pattern distances indicative of increased information content [61]. We assume a correspondence between this information content and the acoustic modulation features used in the neural tracking analysis i.e. enhanced pattern distances associated with stronger neural tracking.

The model’s lexicon (including phonetic transcriptions) was derived from the CELEX database [82]. For the unintelligible context condition, final word prior probabilities in the model’s lexicon were set to normalised word frequencies from CELEX. For final words in strongly predicted and weakly predicted sentences, the prior probabilities were set to the cloze response probabilities provided by Peelle et al. [62]. Any word in the model’s lexicon for which there was no response was set to a small non-zero prior probability. Specifically, we set to 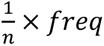 where *n* is the size of the model’s lexicon and *freq* is a weighting factor based on normalised word frequency from CELEX. This adjustment ensured there were no words with a prior probability of zero, which is ill-defined for Bayesian inference [1]. Following this adjustment, prior word probabilities were normalised to sum to one.

The sensory evidence for different words (i.e. likelihoods) was computed by applying the product rule across the individual segment (input) probabilities spanning each word [79]. Segment input probabilities were derived from the acoustic similarity between individual speech segments. This means that an unfolding word’s sensory evidence is strong if its constituent segments match internal representations that are acoustically dissimilar to other speech sounds in the language. It is also strong if there are few word competitors sharing the same speech segments.

To derive segment input probabilities, acoustic representations were first extracted from 4 channel noise-vocoded versions of the sentences, including both context and final words. Acoustic representations were generated by computing a 128 channel ‘auditory’ spectrogram for each segment using the NSL toolbox and averaging over time. Segment onset times were obtained using a forced-alignment algorithm included as part of BAS software (see [83]; https://clarin.phonetik.uni-muenchen.de/BASWebServices/interface/WebMAUSGeneral).

In total there were 44 unique segments, averaged over 8065 repeated tokens (median token count = 142). Acoustic representations were vectorized and averaged over multiple productions of the same segment. We then obtained a between-segment similarity matrix using negative Euclidean distances and normalised each row of this matrix using the softmax function (combined with a temperature parameter) such that each row could be interpreted as segment probabilities. We simulated the three levels of signal quality in Experiment 2 by adjusting the temperature parameter so that the averaged probability of a segment (i.e. the average probability over the on-diagonal elements in the segment matrix) corresponded to .17, .66 and .94. These probabilities simulate the intelligibility levels (proportion correct) observed in previous studies for the three levels of signal quality (4, 8 and 16 channels) used in Experiment 2 (e.g. [84,85]).

## Acknowledgements

This research was supported by a University of Sussex, School of Psychology PhD studentship awarded to JW and University of Sussex, School of Psychology start-up funds awarded to ES. We thank Yiming Zhang and Aine Glynn for assistance with data collection and Chris Racey for technical support.

## Notes

### Competing Interest Statement

The authors have declared no competing interest.

### Summary of Updates

-Minor revisions to Abstract and Introduction -Moved Methods to after Discussion -Changed citation style

## References

1. Davis MH, Sohoglu E. Three functions of prediction error for Bayesian inference in speech perception. In: Poeppel D, Mangun GR, Gazzaniga, Michael S, editors. Cog Neuro. MIT Press; 2020. pp. 177–192.

2. Norris D, McQueen JM, Cutler A. Prediction, Bayesian inference and feedback in speech recognition. Language, Cognition and Neuroscience. 2015;3798: 1–15. doi:10.1080/23273798.2015.1081703

3. Yildiz IB, von Kriegstein K, Kiebel SJ. From Birdsong to Human Speech Recognition: Bayesian Inference on a Hierarchy of Nonlinear Dynamical Systems. PLoS Computational Biology. 2013;9. doi:10.1371/journal.pcbi.1003219

4. McClelland JL, Mirman D, Bolger DJ, Khaitan P. Interactive activation and mutual constraint satisfaction in perception and cognition. Cognitive Science. 2014;38: 1139–1189. doi:10.1111/cogs.12146

5. Gagnepain P, Henson RN, Davis MH. Temporal Predictive Codes for Spoken Words in Auditory Cortex. Current biology : CB. 2012;22: 1–7. doi:10.1016/j.cub.2012.02.015

6. Spratling MW. A review of predictive coding algorithms. Brain and cognition. 2017;112: 92–97. doi:10.1016/j.bandc.2015.11.003

7. Hovsepyan S, Olasagasti I, Giraud A. Combining predictive coding and neural oscillations enables online syllable recognition in natural speech. Nature Communications. 2020;11: 3117. doi:10.1038/s41467-020-16956-5

8. Nour Eddine S, Brothers T, Wang L, Spratling M, Kuperberg GR. A predictive coding model of the N400. Cognition. 2024;246: 105755. doi:10.1016/j.cognition.2024.105755

9. Bastos AMM, Usrey WMM, Adams RAA, Mangun GRR, Fries P, Friston KJJ. Canonical Microcircuits for Predictive Coding. Neuron. 2012;76: 695–711. doi:10.1016/j.neuron.2012.10.038

10. Rao RP, Ballard DH. Predictive coding in the visual cortex: a functional interpretation of some extra-classical receptive-field effects. Nature neuroscience. 1999;2: 79–87. doi:10.1038/4580

11. Kutas M, Federmeier KD. Thirty Years and Counting: Finding Meaning in the N400 Component of the Event-Related Brain Potential (ERP). Annu Rev Psychol. 2011;62: 621–647. doi:10.1146/annurev.psych.093008.131123

12. Lau EF, Phillips C, Poeppel D. A cortical network for semantics: (de)constructing the N400. Nature reviews Neuroscience. 2008;9: 920–33. doi:10.1038/nrn2532

13. Gotts SJ, Chow CC, Martin A. Repetition priming and repetition suppression: Multiple mechanisms in need of testing. Cognitive Neuroscience. 2012;3: 250–259. doi:10.1080/17588928.2012.697054

14. Grill-Spector K, Henson R, Martin A. Repetition and the brain: neural models of stimulus-specific effects. Trends in cognitive sciences. 2006;10: 14–23. doi:10.1016/j.tics.2005.11.006

15. Todorovic A, de Lange FP. Repetition suppression and expectation suppression are dissociable in time in early auditory evoked fields. The Journal of neuroscience : the official journal of the Society for Neuroscience. 2012;32: 13389–95. doi:10.1523/JNEUROSCI.2227-12.2012

16. Blank H, Davis MH. Prediction Errors but Not Sharpened Signals Simulate Multivoxel fMRI Patterns during Speech Perception. PLoS biology. 2016;14: e1002577. doi:10.1371/journal.pbio.1002577

17. De Lange FP, Heilbron M, Kok P. How Do Expectations Shape Perception? Trends in Cognitive Sciences. 2018;22: 764–779. doi:10.1016/j.tics.2018.06.002

18. Luthra S, Li MYC, You H, Brodbeck C, Magnuson JS. Does signal reduction imply predictive coding in models of spoken word recognition? Psychonomic bulletin & review. 2021. doi:10.3758/s13423-021-01924-x

19. Murray SO, Schrater P, Kersten D. Perceptual grouping and the interactions between visual cortical areas. Neural networks : the official journal of the International Neural Network Society. 2004;17: 695–705. doi:10.1016/j.neunet.2004.03.010

20. Donhauser PW, Baillet S. Two Distinct Neural Timescales. Neuron. 2020; 1–9. doi:10.1016/j.neuron.2019.10.019

21. Gillis M, Vanthornhout J, Simon JZ, Francart T, Brodbeck C. Neural Markers of Speech Comprehension: Measuring EEG Tracking of Linguistic Speech Representations, Controlling the Speech Acoustics. J Neurosci. 2021;41: 10316–10329. doi:10.1523/JNEUROSCI.0812-21.2021

22. Heilbron M, Armeni K, Schoffelen J-M, Hagoort P, de Lange FP. A hierarchy of linguistic predictions during natural language comprehension. Proc Natl Acad Sci USA. 2022;119: e2201968119. doi:10.1073/pnas.2201968119

23. Slaats S, Weissbart H, Schoffelen J-M, Meyer AS, Martin AE. Delta-Band Neural Responses to Individual Words Are Modulated by Sentence Processing. J Neurosci. 2023;43: 4867–4883. doi:10.1523/JNEUROSCI.0964-22.2023

24. Weissbart H, Kandylaki KD, Reichenbach T. Cortical Tracking of Surprisal during Continuous Speech Comprehension. Journal of Cognitive Neuroscience. 2020;32: 155–166. doi:10.1162/jocn_a_01467

25. Sohoglu E, Davis MH. Rapid computations of spectrotemporal prediction error support perception of degraded speech. eLife. 2020;9: 1–25. doi:10.7554/eLife.58077

26. Huettig F, Mani N. Is prediction necessary to understand language? Probably not. Language, Cognition and Neuroscience. 2016;31: 19–31. doi:10.1080/23273798.2015.1072223

27. Crosse MJ, Di Liberto GM, Bednar A, Lalor EC, Liberto GMD, Bednar A, et al. The Multivariate Temporal Response Function (mTRF) Toolbox: A MATLAB Toolbox for Relating Neural Signals to Continuous Stimuli. Frontiers in Human Neuroscience. 2016;10: 1–14. doi:10.3389/fnhum.2016.00604

28. Chi T, Ru P, Shamma SA. Multiresolution spectrotemporal analysis of complex sounds. The Journal of the Acoustical Society of America. 2005;118: 887–906. doi:10.1121/1.1945807

29. Loftus GR, Masson MEJ. Using confidence intervals in within-subject designs. Psychonomic Bulletin & Review. 1994;1: 476–490. doi:10.3758/BF03210951

30. Brodbeck C, Simon JZ. Continuous speech processing. Current Opinion in Physiology. 2020;18: 25–31. doi:10.1016/j.cophys.2020.07.014

31. Ahissar E, Nagarajan S, Ahissar M, Protopapas a, Mahncke H, Merzenich MM. Speech comprehension is correlated with temporal response patterns recorded from auditory cortex. Proceedings of the National Academy of Sciences of the United States of America. 2001;98: 13367–72. doi:10.1073/pnas.201400998

32. Luo H, Poeppel D. Phase patterns of neuronal responses reliably discriminate speech in human auditory cortex. Neuron. 2007;54: 1001–10. doi:10.1016/j.neuron.2007.06.004

33. Peelle JE, Gross J, Davis MH. Phase-Locked Responses to Speech in Human Auditory Cortex are Enhanced During Comprehension. Cerebral Cortex. 2012 [cited 21 May 2012]. doi:10.1093/cercor/bhs118

34. Etard O, Reichenbach T. Neural Speech Tracking in the Theta and in the Delta Frequency Band Differentially Encode Clarity and Comprehension of Speech in Noise. The Journal of Neuroscience. 2019;39: 5750–5759. doi:10.1523/JNEUROSCI.1828-18.2019

35. Teoh ES, Ahmed F, Lalor EC. Attention Differentially Affects Acoustic and Phonetic Feature Encoding in a Multispeaker Environment. The Journal of neuroscience : the official journal of the Society for Neuroscience. 2022;42: 682–691. doi:10.1523/JNEUROSCI.1455-20.2021

36. Brodbeck C, Hong LE, Simon JZ. Rapid Transformation from Auditory to Linguistic Representations of Continuous Speech. Current biology : CB. 2018;28: 3976–3983.e5. doi:10.1016/j.cub.2018.10.042

37. Sohoglu E, Beckers L, Davis MH. Convergent neural signatures of speech prediction error are a biological marker for spoken word recognition. Nat Commun. 2024;15: 1–17. doi:10.1038/s41467-024-53782-5

38. Baltzell LS, Srinivasan R, Richards VM. The effect of prior knowledge and intelligibility on the cortical entrainment response to speech. Journal of Neurophysiology. 2017;118: 3144–3151. doi:10.1152/jn.00023.2017

39. Corcoran AW, Perera R, Koroma M, Kouider S, Hohwy J, Andrillon T. Expectations boost the reconstruction of auditory features from electrophysiological responses to noisy speech. Cerebral Cortex. 2022; 2021.09.06.459160. doi:10.1093/cercor/bhac094

40. Di Liberto GM, Crosse MJ, Lalor EC. Cortical Measures of Phoneme-Level Speech Encoding Correlate with the Perceived Clarity of Natural Speech. eNeuro. 2018;5: ENEURO.0084-18.2018. doi:10.1523/ENEURO.0084-18.2018

41. Holdgraf CR, de Heer W, Pasley B, Rieger J, Crone N, Lin JJ, et al. Rapid tuning shifts in human auditory cortex enhance speech intelligibility. Nature Communications. 2016;7: 13654. doi:10.1038/ncomms13654

42. Karunathilake IMD, Kulasingham JP, Simon JZ. Neural tracking measures of speech intelligibility: Manipulating intelligibility while keeping acoustics unchanged. Proc Natl Acad Sci USA. 2023;120: e2309166120. doi:10.1073/pnas.2309166120

43. Molinaro N, Lizarazu M, Baldin V, Pérez-Navarro J, Lallier M, Ríos-López P. Speech-brain phase coupling is enhanced in low contextual semantic predictability conditions. Neuropsychologia. 2021;156: 107830. doi:10.1016/j.neuropsychologia.2021.107830

44. Davis MH, Johnsrude IS, Hervais-Adelman A, Taylor K, McGettigan C. Lexical information drives perceptual learning of distorted speech: evidence from the comprehension of noise-vocoded sentences. Journal of experimental psychology General. 2005;134: 222–41. doi:10.1037/0096-3445.134.2.222

45. Kok P, Jehee JFM, de Lange FP. Less is more: expectation sharpens representations in the primary visual cortex. Neuron. 2012;75: 265–70. doi:10.1016/j.neuron.2012.04.034

46. Obleser J, Kotz S a. Expectancy constraints in degraded speech modulate the language comprehension network. Cerebral cortex (New York, NY : 1991). 2010;20: 633–40. doi:10.1093/cercor/bhp128

47. Obleser J, Kotz SA. Multiple brain signatures of integration in the comprehension of degraded speech. NeuroImage. 2011;55: 713–23. doi:10.1016/j.neuroimage.2010.12.020

48. Holdgraf CR, Rieger JW, Micheli C, Martin S, Knight RT, Theunissen FE. Encoding and Decoding Models in Cognitive Electrophysiology. Frontiers in systems neuroscience. 2017;11: 61. doi:10.3389/fnsys.2017.00061

49. Alink A, Schwiedrzik CM, Kohler A, Singer W, Muckli L. Stimulus predictability reduces responses in primary visual cortex. The Journal of neuroscience : the official journal of the Society for Neuroscience. 2010;30: 2960–6. doi:10.1523/JNEUROSCI.3730-10.2010

50. Murray SO, Kersten D, Olshausen B a, Schrater P, Woods DL. Shape perception reduces activity in human primary visual cortex. Proceedings of the National Academy of Sciences of the United States of America. 2002;99: 15164–9. doi:10.1073/pnas.192579399

51. Thomas ER, Haarsma J, Nicholson J, Yon D, Kok P, Press C. Predictions and errors are distinctly represented across V1 layers. Current Biology. 2024;34: 2265–2271.e4. doi:10.1016/j.cub.2024.04.036

52. Sohoglu E, Davis MH. Perceptual learning of degraded speech by minimizing prediction error. Proceedings of the National Academy of Sciences. 2016;113: E1747–E1756. doi:10.1073/pnas.1523266113

53. Widmann A, Schröger E, Maess B. Digital filter design for electrophysiological data – a practical approach. Journal of Neuroscience Methods. 2015;250: 34–46. doi:10.1016/j.jneumeth.2014.08.002

54. Klimovich-Gray A, Barrena A, Agirre E, Molinaro N. One Way or Another: Cortical Language Areas Flexibly Adapt Processing Strategies to Perceptual And Contextual Properties of Speech. Cerebral cortex (New York, NY : 1991). 2021;31: 4092–4103. doi:10.1093/cercor/bhab071

55. Mai G, Wang WS-Y. Distinct roles of delta- and theta-band neural tracking for sharpening and predictive coding of multi-level speech features during spoken language processing. Human Brain Mapping. 2023;44: 6149–6172. doi:10.1002/hbm.26503

56. Marslen-Wilson W. The temporal structure of spoken language understanding. Cognition. 1980;8: 1–71. doi:10.1016/0010-0277(80)90015-3

57. McClelland JL, Elman JL. The TRACE model of speech perception. Cognitive Psychology. 1986;18: 1–86. doi:10.1016/0010-0285(86)90015-0

58. Norris D. Shortlist: A connectionist model of continuous speech recognition. Cognition. 1994;52: 189–234.

59. Spratling MW. A neural implementation of Bayesian inference based on predictive coding. Connection Science. 2016;28: 346–383. doi:10.1080/09540091.2016.1243655

60. Millidge B, Tang M, Osanlouy M, Harper NS, Bogacz R. Predictive coding networks for temporal prediction. PLOS Computational Biology. 2024;20: e1011183. doi:10.1371/journal.pcbi.1011183

61. Kriegeskorte N, Kievit RA. Representational geometry: integrating cognition, computation, and the brain. Trends in Cognitive Sciences. 2013;17: 401–412. doi:10.1016/j.tics.2013.06.007

62. Peelle JE, Miller RL, Rogers CS, Spehar B, Sommers MS, Van Engen KJ. Completion norms for 3085 English sentence contexts. Behavior research methods. 2020;52: 1795–1799. doi:10.3758/s13428-020-01351-1

63. Taylor WL. “Cloze Procedure”: A New Tool for Measuring Readability. Journalism Quarterly. 1953;30: 415–433. doi:10.1177/107769905303000401

64. Bloom PA, Fischler I. Completion norms for 329 sentence contexts. Memory & Cognition. 1980;8: 631–642. doi:10.3758/BF03213783

65. Gwilliams L, Davis M. Extracting language content from speech sounds: An information theoretic approach. In: Holt LL, Peelle JE, Coffin AB, Popper AN, Fay RR, editors. Speech Perception. Springer; 2022. pp. 113–139.

66. Shannon RV, Zeng F-G, Kamath V, Wygonski J, Ekelid M. Speech Recognition with Primarily Temporal Cues. Science. 1995;270: 303–304. doi:10.1126/science.270.5234.303

67. Greenwood DD. A cochlear frequency-position function for several species—29 years later. Journal of the Acoustical Society of America. 1990;87: 2592–2605. doi:10.1121/1.399052

68. de Cheveigné A, Arzounian D. Robust detrending, rereferencing, outlier detection, and inpainting for multichannel data. NeuroImage. 2018;172: 903–912. doi:10.1016/j.neuroimage.2018.01.035

69. de Cheveigné A. ZapLine: A simple and effective method to remove power line artifacts. NeuroImage. 2020;207: 116356. doi:10.1016/j.neuroimage.2019.116356

70. Di Liberto GM, O’Sullivan JA, Lalor EC. Low-frequency cortical entrainment to speech reflects phoneme-level processing. Current Biology. 2015;25: 2457–2465. doi:10.1016/j.cub.2015.08.030

71. Ding N, Simon JZ. Adaptive temporal encoding leads to a background-insensitive cortical representation of speech. The Journal of neuroscience : the official journal of the Society for Neuroscience. 2013;33: 5728–35. doi:10.1523/JNEUROSCI.5297-12.2013

72. Pasley BN, David SV, Mesgarani N, Flinker A, Shamma S a, Crone NE, et al. Reconstructing speech from human auditory cortex. PLoS biology. 2012;10: e1001251. doi:10.1371/journal.pbio.1001251

73. Theunissen FE, Elie JE. Neural processing of natural sounds. Nature reviews Neuroscience. 2014;15: 355–66. doi:10.1038/nrn3731

74. Elliott TM, Theunissen FE. The modulation transfer function for speech intelligibility. PLoS computational biology. 2009;5: e1000302. doi:10.1371/journal.pcbi.1000302

75. Henson RN, Wakeman DG, Litvak V, Friston KJ. A Parametric Empirical Bayesian Framework for the EEG/MEG Inverse Problem: Generative Models for Multi-Subject and Multi-Modal Integration. Frontiers in Human Neuroscience. 2011;5: 1–16. doi:10.3389/fnhum.2011.00076

76. Litvak V, Friston K. Electromagnetic source reconstruction for group studies. NeuroImage. 2008;42: 1490–8. doi:10.1016/j.neuroimage.2008.06.022

77. Phillips C, Mattout J, Rugg MD, Maquet P, Friston KJ. An empirical Bayesian solution to the source reconstruction problem in EEG. NeuroImage. 2005;24: 997–1011. doi:10.1016/j.neuroimage.2004.10.030

78. Friston K, Henson R, Phillips C, Mattout J. Bayesian estimation of evoked and induced responses. Human brain mapping. 2006;27: 722–35. doi:10.1002/hbm.20214

79. Norris D, McQueen JM. Shortlist B: a Bayesian model of continuous speech recognition. Psychological review. 2008;115: 357–95. doi:10.1037/0033-295X.115.2.357

80. Aitchison L, Lengyel M. With or without you: predictive coding and Bayesian inference in the brain. Current Opinion in Neurobiology. 2017;46: 219–227. doi:10.1016/j.conb.2017.08.010

81. Brodbeck C, Bhattasali S, Cruz Heredia AA, Resnik P, Simon JZ, Lau E. Parallel processing in speech perception with local and global representations of linguistic context. eLife. 2022;11: e72056. doi:10.7554/eLife.72056

82. Baayen RH, Piepenbrock R, Van Rijn H. The CELEX lexical database (CD-ROM). Philadelphia Linguistics Data Consortium University of Pennsylvania. 1993.

83. Kisler T, Reichel U, Schiel F. Multilingual processing of speech via web services. Computer Speech & Language. 2017;45: 326–347. doi:10.1016/j.csl.2017.01.005

84. McGettigan C, Rosen S, Scott SK. Lexico-semantic and acoustic-phonetic processes in the perception of noise-vocoded speech: implications for cochlear implantation. Frontiers in systems neuroscience. 2014;8: 18. doi:10.3389/fnsys.2014.00018

85. Sohoglu E, Peelle JE, Carlyon RP, Davis MH. Predictive Top-Down Integration of Prior Knowledge during Speech Perception. The Journal of neuroscience : the official journal of the Society for Neuroscience. 2012;32: 8443–8453. doi:10.1523/JNEUROSCI.5069-11.2012

